# HGCF/LGCF classification from (Mavrich and Hatfull, 2017) is explained by heterogeneity in pairwise ANI rather than by intrinsic evolutionary modes

**DOI:** 10.64898/2026.11.23.768377

**Authors:** Alexandre Boulay, Clovis Galiez

**Affiliations:** Univ. Grenoble Alpes, CNRS, Grenoble INP, LJK, 38000 Grenoble, France

## Abstract

Mavrich and Hatfull proposed that phages could be separated into two broad evolutionary modes (high gene-content flux or HGCF; and low gene-content flux or LGCF) according to the rate at which their gene content changes relative to nucleotide sequence divergence. In their analysis, HGCF phages appeared to gain and lose genes substantially faster relative to genome divergence than LGCF phages and were particularly associated with temperate phages. However, as the analysis represents a comparison between two phage genomes, the difference between the two groups identified in their study (HGCF/LGCF) should not be interpreted as a property of either phage (such as their lifestyle) as they suggest. Instead, it should be explained by a property inherent to the comparison such as the organization of the shared regions between both genomes. Indeed, the two groups identified in the original study reflect differences in the ANI (average nucleotide identity) and coverage of aligned regions between pairs of genomes and is independent from the gene-content similarity central to the original interpretation. More fundamentally, we show that the k-mer based genomic similarity used in the original study (namely Mash) is biased towards short regions of very high nucleotide identity, which exaggerates the differences described previously. This is particularly important for phage genomes which are often mosaic: closely related regions can alternate with more divergent or unrelated regions, and the number, length and identity of these regions can differ markedly between genome pairs.

## Background context

Mavrich and Hatfull proposed that phages could be separated into two broad evolutionary modes (high gene-content flux or HGCF; and low gene-content flux or LGCF) according to the rate at which their gene content changes relative to nucleotide sequence divergence (Mavrich and Hatfull, 2017) (**Figure 1A**). In their analysis, HGCF phages appeared to gain and lose genes substantially faster than LGCF phages and were particularly associated with temperate phages. Moura de Sousa et al. subsequently provided a potential mechanistic explanation for this observation. Temperate HGCF phages were considerably more likely to encode recombinases than temperate LGCF phages (62% versus 27%) and were thus proposed as systems that could underlie their higher gene flux (Moura de Sousa et al., 2021). These observations supported the interpretation that some phages recombine more frequently and consequently undergo more rapid gene-content turnover.

**Figure 1.**
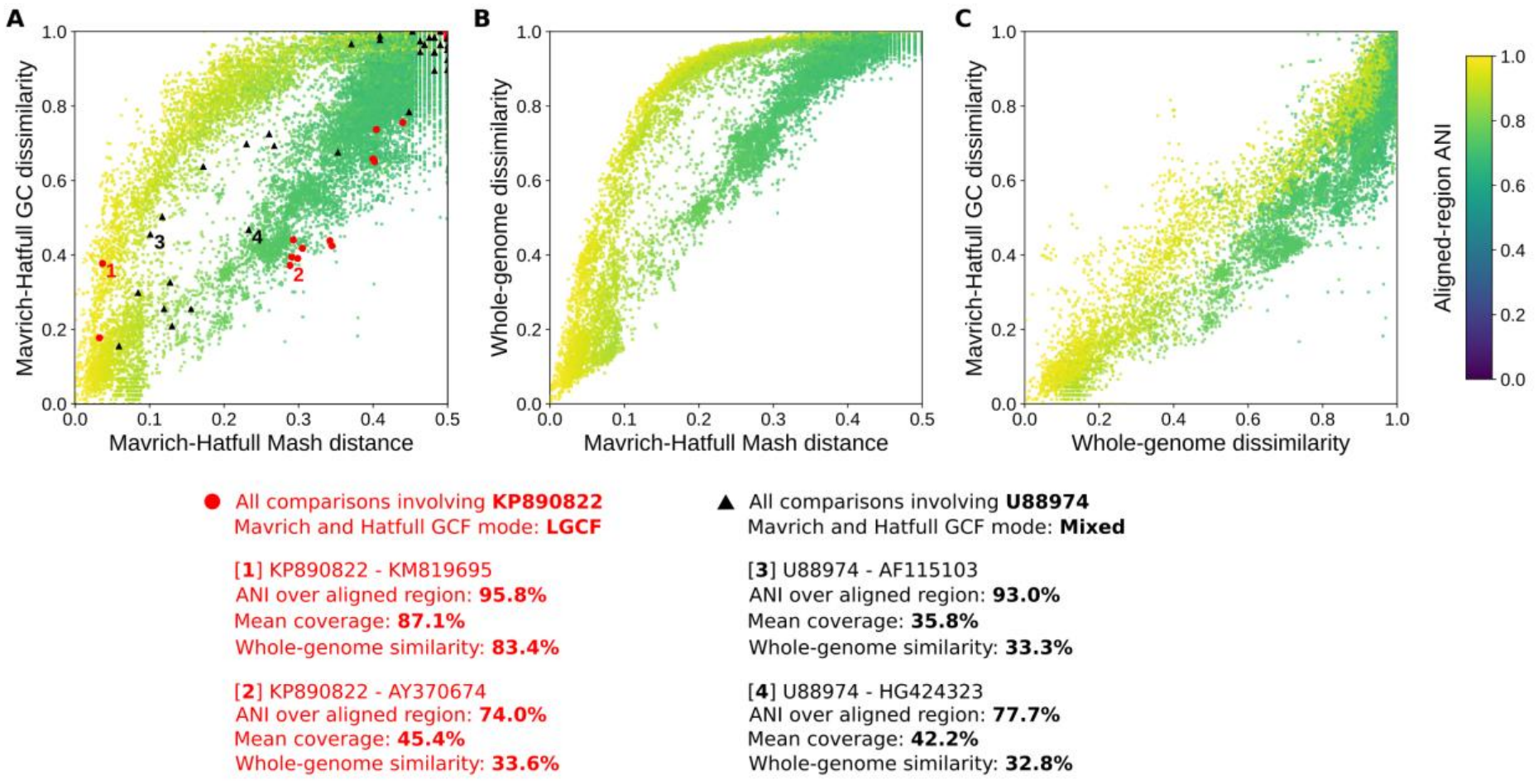
Similarity between 30,080 pairs of phages from the original Mavrich and Hatfull study (2017). **A)** Relationship between the nucleotide dissimilarity estimated using Mash and the gene content dissimilarity of pham clusters between phages being compared (from Mavrich and Hatfull). The red circles and black triangles highlight all comparisons involving KP890822 and U88974, respectively. Additional alignment details are provided for an “HGCF” and an “LGCF” comparison involving each phage. **B)** Relationship between the whole-genome nucleotide distance calculated using MMseqs2 and the Mash nucleotide distance from Mavrich and Hatfull. **C)** Relationship between the whole-genome nucleotide distance calculated using MMseqs2 and the gene content dissimilarity of pham clusters between phages being compared. All figures are coloured using the aligned-region ANI (Average Nucleotide Identity) measured using MMseqs2.

## Methodological and analytical limitations of the HGCF/LGCF comparison

Firstly, as each point in the HGCF/LGCF analysis represents a comparison between two genomes, the difference between the two curves should not be interpreted directly as a property of either phage, or of its lifestyle (**Figure 1A**). Instead, it should be explained by a property inherent to the comparison *between both genomes* such as the organization of the shared regions between them. This is particularly important for phage genomes which are often mosaic: closely related regions can alternate with more divergent or unrelated regions, and the number, length and identity of these regions can differ markedly between genome pairs (Casjens, 2003; Belcaid et al., 2010; Pope et al., 2015).

Secondly, as observed in **Figure 1B**, the same “distinct” (HGCF/LGCF) groups can be observed in a direct comparison between the nucleotide dissimilarity estimated by Mash and the one calculated using MMseqs2 in nucleotide mode (as described below), despite neither measure involving protein-content similarity. Consistently, gene content dissimilarity measured by Mavrich and Hatfull is more linearly associated with whole genome dissimilarity than with Mash distance (**Figure 1C**). These figures suggest that the relationship between HGCF/LGCF phages is not directly a result of phages exchanging proteins at a higher or lower rate and that the measure estimated by Mash is misleading in this regard. As suggested previously, these observations could be explained by a difference in the genome organization and nucleotide similarity between pairs of HGCF and LGCF phages.

In particular, differences in the organization of shared sequence – for example, whether similarity is concentrated in a short region with very high identity or distributed across a longer fraction of both genomes – could play a role in the nucleotide distance reported by Mash (Ondov et al., 2016). This can already be observed in the colour displayed in **Figure 1** which shows ANI (Average Nucleotide Identity) over only the aligned region of genome pairs. As further demonstrated in **Supplementary Figure 1**, the distance estimated by Mash does not follow a linear relationship with either ANI or alignment coverage.

Along these lines, the nucleotide distance calculated by Mash is an estimate of sequence divergence under a simple evolutionary model that is based on the fraction of shared k-mers between two genomes. Mash distance increases as the fraction of shared k-mers decreases (**Figure 2; Supplementary Note 1**). However, this relationship is nonlinear and phage mosaicity further complicates the biological interpretation of the Mash distance. Two phage genomes can simultaneously contain nearly identical regions, regions that remain similar but have accumulated substantial sequence divergence, and regions with little or no detectable nucleotide similarity. Each of these regions contributes differently to the genome-wide fraction of shared k-mers. Nearly identical regions contribute large numbers of shared k-mers and keep Mash distance close to zero, whereas increasingly diverged regions contribute progressively fewer exact k-mers. Because of the nonlinear transformation used by Mash, these contributions are not reflected proportionally in the final distance estimate. Consequently, relatively small genomic regions with very high nucleotide identity can contribute disproportionately to the shared k-mer fraction and pull the estimated Mash distance toward zero.

**Figure 2.**
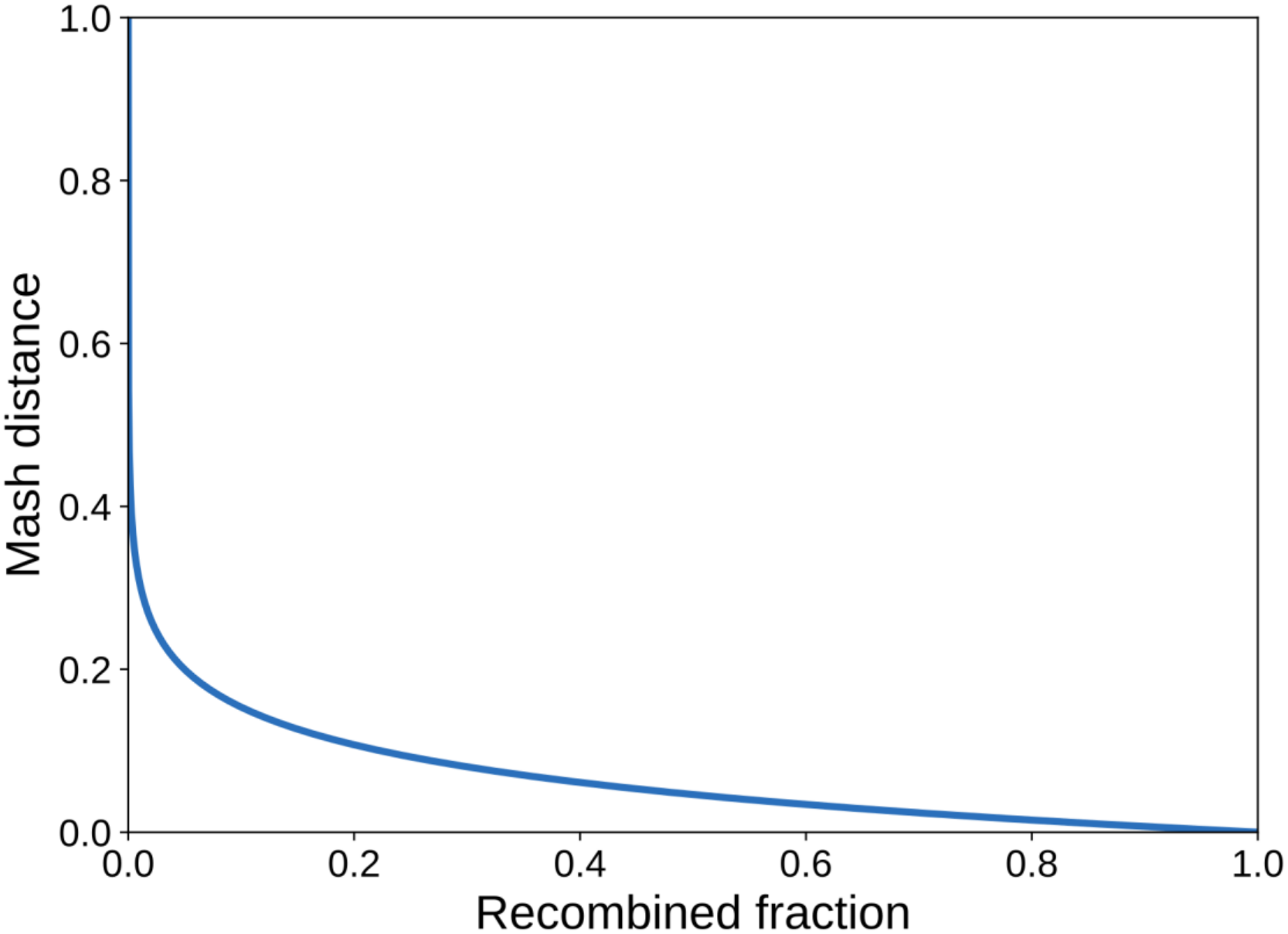
Relationship between Mash distance and the fraction of two otherwise unrelated genomes sharing an identical sequence (using a kmer size of 15).

## Reinterpreting HGCF and LGCF curves

An important issue with the original interpretation is the assumption that “LGCF” phages undergo less gene exchange than “HGCF” phages. Moura de Sousa et al. (2021) already challenged this idea: “Even if we observe four times more recent transfers between temperate phages than between other pairs, there is extensive gene flow between temperate and […] virulent phages with large genomes previously classed as low gene flux”. Indeed, gene flux should theoretically impact both the gene content and nucleotide similarities of *pairwise comparisons* relatively proportionally: the acquisition/loss of a protein will also necessarily alter the similarity between their genomes in a similar way. The distinction between “HGCF” and “LGCF” should therefore not be interpreted as a direct measure of how much genetic exchange a phage lineage undergoes.

Rather than corresponding to a rate of protein exchange relative to nucleotide sequence divergence, as proposed by Mavrich and Hatfull (2017), the two “HGCF/LGCF” curves correspond to pairs of phages sharing different kinds of genomic similarities. The bottom curve, previously characterized as “LGCF”, corresponds to pairs of phages that share common genomic regions, but that have diverged over time (i.e. a lower ANI). The “HGCF” curve, for its part, corresponds to phages sharing a very high ANI, potentially a result of a more recent recombination between two distinct phages, or of the acquisition/exchange of new genetic material in at least one of two very closely related phages. This new interpretation also explains why a same phage can be found in both curves depending on the phage it is being compared to (**Figure 1A**), a pattern that had required Mavrich and Hatfull to introduce an ambiguous “Mixed” category in their original classification.

While it may be true that “HGCF” phages have exchanged genetic material more recently with the phage(s) it is being compared to, this is by no means a generalizable trait of the *propensity* for genetic exchange characteristic of the phage in nature. For example, a single recombination event between two phages, followed by extensive sampling of descendants from one of these lineages, could generate many such “HGCF” pairs without being the result of elevated gene flux. Furthermore, the prevalence of these pairs would be very dependent on the size and diversity of the sampled dataset.

Other studies have encountered problems when attempting to reconcile their observations with the original HGCF/LGCF interpretation, which they attempt to explain in roundabout ways (Accetto & Janež, 2018; Lavelle et al., 2018; Forni et al., 2024). Moreover, the HGCF/LGCF classification itself has been reused directly, including in the highly impactful vConTACT2 paper and in the analysis of genetic exchange by Moura de Sousa et al. (Bin Jang et al., 2019; Moura de Sousa et al., 2021). Finally, several studies have cited the broader conclusion that phages differ in their rates of protein exchange (Sharma et al., 2023; Kobakhidze et al., 2024; Zhou et al., 2026; Leprince et al., 2026). These papers show that it is important to reassess the basis of the original interpretation as it has continued to influence phage classification and the framing of evolutionary analyses.

### Methods: measuring pairwise genome similarities with MMseqs

MMseqs2 (Steinegger and Söding, 2017) was used to compute the pairwise similarity between all phage genomes present in the analysis. To account for circular genomes, sequences in the target database were doubled (sequences copied back-to-back). Then the mmseqs search command (with --search-type 3 -s 7 --max-seqs 40000 --max-seq-len 10000 - -sequence-overlap 8000) was used. These last 2 parameters were used to more sensitively account for gaps and rearrangements often observed in phage genomes.

These settings, necessary for their increased sensitivity, lead to many small alignments (often duplicates) being reported. A consolidation step was then applied to reconstruct genome-wide similarity from these small local alignments while making sure to account for the overlap between alignments. For each query-target pair, HSPs (High-scoring Segment Pairs) were ranked by bit-score and only query regions not already covered by a higher scoring HSP were considered in the computation of the final score. Because only the alignment statistics (coverage, identity, etc.) were available for each HSP, the number of identical bases was estimated from the nucleotide identity of each contributing HSP segment. For each non-redundant segment of an HSP, the number of identical bases to add to the global-alignment statistics was calculated as the number of new (not already counted) bases multiplied by the measured identity of the whole HSP. Finally, the query similarity was calculated as the estimated number of identical bases divided by the length of the query genome.

For each pair of phages, the directional genome similarities were averaged to obtain a symmetric nucleotide similarity that accounts for differences in genome lengths and aligned coverage.

Only phages from Mavrich and Hatfull’s original analysis for which NCBI accessions were easily retrievable and present in the INPHARED dataset (Cook et al. 2021) were considered in the analysis. A total of 30,080 phage comparisons were retained for the plots shown in **Figure 1**.

## Acknowledgements

The computations presented in this paper were performed using the GRICAD infrastructure (https://gricad.univ-grenoble-alpes.fr), which is supported by Grenoble research communities.

## Supplementaries

### Supplementary Note 1: Derivation of the equivalent mutation rate

Mash uses the following distance formula:

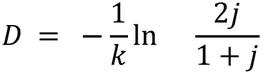

Where j is the Jaccard index of the set of k-mers of each genome. When unrelated genomes of similar length G recombine over a fraction *α* of their genome, the Jaccard index of their set of k-mers is tightly lower bounded by 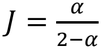 . In that case, the Mash distance amounts to 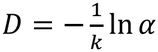 ln *α*. Plotting this distance with respect to *α* (Figure 2) shows that the Mash distance decreases quickly for small recombination ratios.

## Supplementary Figure 1

**Supplementary Figure 1.**
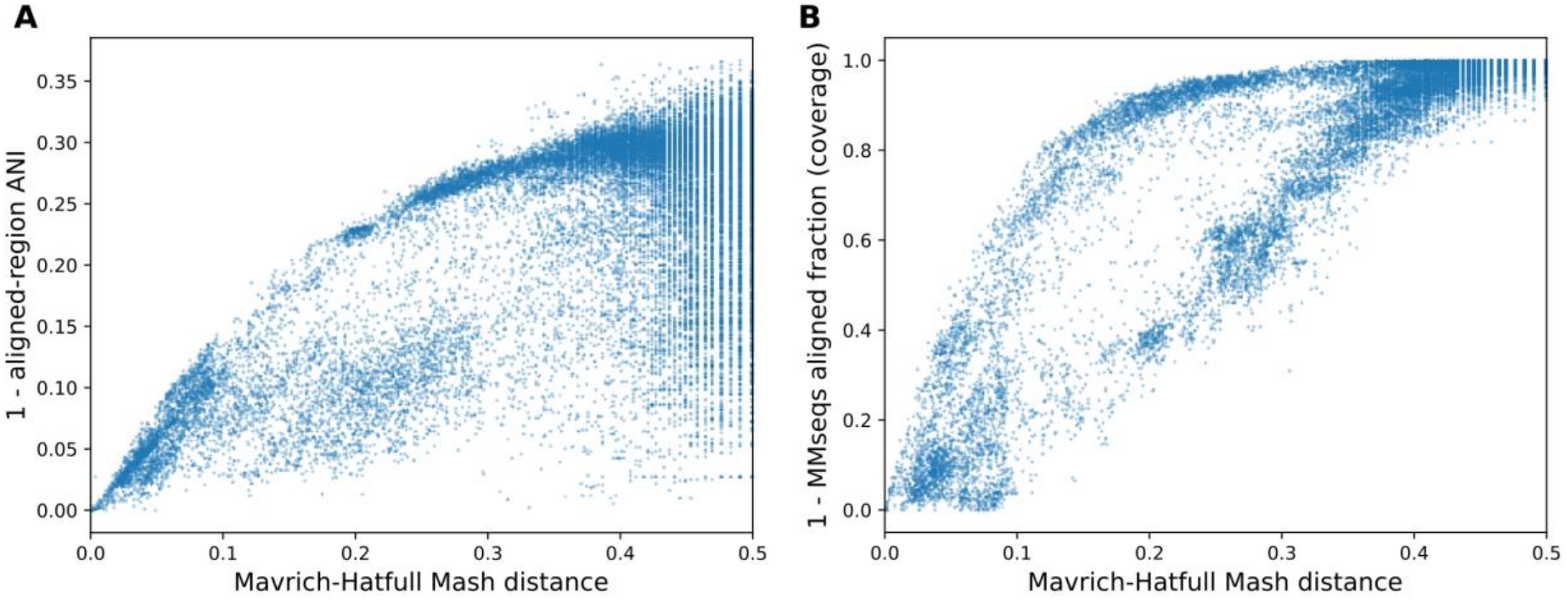
Similarity between 30,080 pairs of phages from the original Mavrich and Hatfull study (2017). **A)** Relationship between the nucleotide dissimilarity estimated using Mash and the ANI over aligned regions measured using MMseqs for the phages being compared. **B)** Relationship between the nucleotide dissimilarity estimated using Mash and the aligned fraction (coverage) measured using MMseqs for the phages being compared.

## Notes

### Competing Interest Statement

The authors have declared no competing interest.

